# Characterization of the nanomechanical properties of the fission yeast (*Schizosaccharomyces pombe*) cell surface by atomic force microscopy

**DOI:** 10.1101/2021.01.18.427162

**Authors:** Ellie Gibbs, Justine Hsu, Kathryn Barth, John W. Goss

## Abstract

Variations in cell wall composition and biomechanical properties can contribute to the cellular plasticity required during complex processes such as polarized growth and elongation in microbial cells. This study utilizes atomic force microscopy (AFM) to map the cell surface topography of fission yeast, *Schizosaccharomyces pombe*, at regions of active polarized growth and to characterize the biophysical properties within these regions under physiological, hydrated conditions. High-resolution images acquired from AFM topographic scanning reveal decreased surface roughness at actively growing cell poles. Force extension curves acquired by nanoindentation probing with AFM cantilever tips under low applied force revealed increased cell wall elasticity and decreased cellular stiffness (cellular spring constant) at cell poles (17 ± 4 mN/m) relative to the main body of the cell that is not undergoing growth and expansion (44 ± 10 mN/m). These findings suggest that the increased elasticity and decreased stiffness at regions undergoing polarized growth at fission yeast cell poles provide the plasticity necessary for cellular extension. This is the first direct biophysical characterization of the *S. pombe* cell surface by AFM, and it provides a foundation for future investigation of how the surface topography and local nanomechanical properties vary during different cellular processes.

## INTRODUCTION

Fungal cells are surrounded by an exterior cell wall that provides shape and structure, resistance to internal turgor pressure, and protection from extracellular physical and environmental stresses. Many mechanisms and proteins regulating synthesis and remodeling of the fission yeast, *Schizosaccharomyces pombe*, cell wall have been characterized through genetic, biochemical, and microscopic studies, but little is known about the *S. pombe* cell wall biophysical properties. The fission yeast cell wall is composed of three major layers: an outer layer of galactomannan, a middle layer of α-(1,3)-glucans, β-(1,3)-glucans, and β-(1,6)-glucans, and an inner layer of galactomannan adjacent to the plasma membrane (M. Horisberger, Vonlanthen, & Rosset, 1978; Humbel et al., 2001; Kopecka, Fleet, & Phaff, 1995; Sugawara, Takahashi, Osumi, & Ohno, 2004). Despite being firm, providing support, and enabling resistance to forces (not unlike the extracellular matrix of animal cells), the cell wall must also be capable of plasticity and remodeling to enable cellular elongation during polarized growth or septum formation during cell division. Furthermore, biochemical studies reveal that the fission yeast cell wall glucan composition can differ in certain regions of the cell or throughout the cell cycle (Cortes et al., 2012; M Horisberger & Rouvet-Vauthey, 1985; M. Horisberger et al., 1978; Humbel et al., 2001). This variation in composition at the molecular level could contribute to differences in the cellular biophysical properties, though this has not been directly evaluated. *S. pombe* is a widely studied model organism for polarized growth, which requires the coordination and integration of cellular signaling pathways, membrane trafficking and vesicle fusion, cytoskeletal organization, and *de novo* cell wall synthesis (Bendezu & Martin, 2011; Cortes et al., 2005; Feierbach & Chang, 2001; Martin & Arkowitz, 2014; Martin, McDonald, Yates, & Chang, 2005; Rincon, Estravis, & Perez, 2014; Sawin & Nurse, 1998). Experimental tools that characterize these biophysical components and their spatial and temporal fluctuations can provide valuable insight into the complex mechanisms that regulate polarized cellular growth.

Atomic force microscopy (AFM) is a scanning probe spectroscopy technique that utilizes a cantilever tip under near-physiological conditions to map surfaces at nanoscale resolution and investigate their mechanical properties through nanoindentation of the cellular exterior (Alessandrini & Facci, 2005; Dufrene, 2002; Dufrene et al., 2017; Goss & Volle, 2019; Hansma et al., 1994; Horber & Miles, 2003). The pressure of the cantilever tip upon the cell surface under low applied force yields a force vs. indentation curve that can provide a wealth of information about cellular biophysical parameters (Burks et al., 2003; Pelling, Sehati, Gralla, Valentine, & Gimzewski, 2004; Velegol & Logan, 2002; Volle, Ferguson, Aidala, Spain, & Nunez, 2008). A variety of indirect measurements or computational approaches have previously been implemented to model the mechanical properties of fission yeast cells (Abenza et al., 2015; Atilgan, Magidson, Khodjakov, & Chang, 2015; Davi et al., 2018; Minc, Boudaoud, & Chang, 2009), but prior utilization of AFM in *S. pombe* has not expanded beyond mapping cell surface topography or evaluating ligand-receptor adhesion (Adya, Canetta, & Walker, 2006; Canetta, Adya, & Walker, 2006; Canetta, Walker, & Adya, 2009; Ishijima et al., 1999; Sasuga et al., 2012). To date, no direct quantification of *S. pombe* cell wall biophysical properties have been experimentally measured. AFM has been employed much more widely in the direct biophysical characterization of budding yeast, *Saccharomyces cerevisiae*, cell wall mechanical properties such as stiffness, elasticity, turgor pressure, and thickness leading to an enhanced understanding of how the cell wall and cellular turgor pressure differ during a variety of cellular processes and in response to environmental stressors (Alsteens et al., 2008; Dague et al., 2010a; Dupres, Dufrene, & Heinisch, 2010; Pelling et al., 2004; A. Touhami, Nysten, & Dufrene, 2003).

This study utilizes AFM contact mode scanning to generate high resolution images of the *S. pombe* cell surface topography and force extension curves generated from nanoindentation to quantify the cell wall elasticity and cellular spring constant. These biophysical components were then evaluated at the fission yeast cell body or cell pole to determine whether regions undergoing polarized growth have different mechanical properties. Regions of cellular extension at cell poles had decreased surface roughness, increased cell wall elasticity, and decreased cellular stiffness relative to the cell body where no active growth or expansion occurs. These findings provide the first nanomechanical characterization of fission yeast by AFM that serves as the foundation for future scanning probe spectroscopic studies to advance the analyses of polarized growth, cell wall biogenesis, and cellular response to environmental changes.

## MATERIALS AND METHODS

### Preparation of dishes for cell adhesion

Plastic 50 × 9 mm petri dishes (Corning Inc., Corning, NY) were prepared for yeast cell adhesion by plasma cleaning for 3 minutes with PDC-32G plasma cleaner (Harrick Plasma, Ithaca, NY). Plates were then incubated for 48 hours at 4°C with 0.8 mg/mL CellTak adhesive (Corning Life Sciences, Glendale, AZ) diluted in 0.1M NaHCO_3_. Treated dishes were washed twice with ddH_2_O to remove excess CellTak and allowed to air dry for 5 minutes prior to plating cells.

### Strains, growth conditions, and plating fission yeast cells for atomic force microscopy

*Schizosaccharomyces pombe* strains used in this study are provided in Supplemental Table 1. Cells were cultured in YE5S medium at 25°C in mid-log phase (OD_595_ <0.6) for 36 hours on a rotary wheel prior to microscopic studies. Cells for AFM imaging and analysis were centrifuged at 500 x g for 2 minutes and washed twice with EMM5S growth media prior to plating on CellTak treated plastic dishes. 200 μL suspension of washed cells was added to treated dish and centrifuged at 500 x g for 20 seconds using a swinging bucket rotor. Dishes were washed with EMM5S three times to remove non-adherent cells, and remaining cells were incubated in 100 μL EMM5S for imaging. Preliminary experiments testing different AFM buffers also utilized HEPES (0.1 mM HEPES, 0.01 mM CaCl_2_, 0.001 mM MgCl_2_, pH 7.0) or sodium acetate (18 mM sodium acetate, 1 mM CaCl_2_, 1 mM MnCl_2_, pH 5.2) buffers. All imaging experiments were conducted within 90 minutes of adhering cells.

For imaging and analysis of cell poles by AFM, overnight liquid cultured cells were centrifuged at 500 x g for 2 minutes and washed twice with EMM5S medium. A 5 μm pore diameter polycarbonate isopore membrane filter (22 μm thickness) (Millipore, Burlington, MA) was stacked on top of a 0.2 μm membrane (Pall Corp, New York, NY) and both membranes were placed in a polysulfone bottle top filter (Thermo Fisher, Waltham, MA). The cell suspension was added to the membranes and low vacuum suction was briefly applied to draw cells into membrane wells. Membranes were removed and excess cells were washed away with EMM5S. The membrane containing cells was cut into 2 cm x 2 cm squares, adhered to a glass slide using double-sided tape, further secured to the slide using paraffin wax, and incubated in 100 μL EMM5S media for imaging.

### Atomic force microscopy imaging and analysis

Topographical height and deflection images were obtained in scanning contact mode using an MFP-3D AFM (Asylum Research, Santa Barbara, CA). Silicon Nitride AFM pyramidal tip PNP-TR probes (Nanoworld, Neuchatel, Switzerland) with a nominal tip radius of <10nm and nominal spring constant of 0.32 N/m were used for all experiments. The cantilever spring constant (*k*_cantilever_) was experimentally calculated using the thermal tuning method (Hutter & Bechhoefer, 1993) prior to each experiment through the MFP-3D software. Cells were incubated in EMM5S media throughout AFM imaging and analysis to prevent dehydration. 2D images were exported from MFP-3D software and scale bars were added using NIH Image J. Following topographical imaging, force measurements were obtained from yeast cells at designated locations along the apex of curvature of the cell body or at the cell pole at a deflection set point of 4 nN to minimize damage to the cell surface. Spring constants (*k*_cell_) were calculated from the linear region of force extension curves using a two-spring model with the equation:

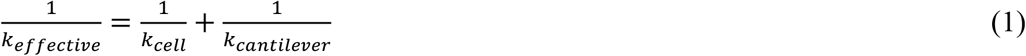

where the slope of the linear region is the *k*_effective_, and *k*_cantilever_ was determined through cantilever calibration (Burks et al., 2003; Pelling et al., 2004; Velegol & Logan, 2002; Volle et al., 2008). The non-linear region of force extension curves was analyzed by separately measuring the change in force and distance. Data processing and analysis for both regions of the extension force curves was performed using the Asylum Igor Pro MFP-3D software. Measurement of surface peak height and roughness was determined from 2D exported height retrace images using NIH Image J with plugin SurfCharJ 1q to calculate root mean square values (Chinga, Johnsen, Dougherty, Berli, & Walter, 2007).

Young’s elasticity moduli were calculated from AFM extension curves by the Asylum Igor Pro MFP-3D software using the Hertz model equation (Ahmed Touhami, Nysten, & Dufrêne, 2003; Zemła et al., 2020):

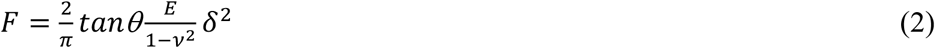

Where *F* is the applied force, δ is the resulting indentation depth, *E* is the Young’s modulus, and *θ* is the half-opening angle (35° from manufacturer). Notably, we utilized a sample Poisson of 0.03 in these calculations as this value was utilized in prior *S. pombe* elasticity modeling (Abenza et al., 2015; Atilgan et al., 2015; Davi et al., 2018). Modeling with the Poisson value of 0.5 commonly used for soft biological materials (Guz, Dokukin, Kalaparthi, & Sokolov, 2014; Ahmed Touhami et al., 2003) resulted in 25% lower Young’s moduli (Supplemental Figure 2).

## RESULTS AND DISCUSSION

### Visualization of fission yeast cells by topography mapping

AFM contact mode scanning was used to evaluate the high-resolution topography of wildtype *S. pombe* cells adhered to plastic dishes. All AFM scanning and analysis was conducted in EMM5S fission yeast minimal growth media to maintain cell hydration and ensure cell viability. No differences in the biophysical properties of fission yeast were observed based on media used during imaging (Supplemental Figure 1). At scan sizes of 10 μm, deflection retrace images from contact mode scanning along the surface of cells (at <1 nN applied force) reveal the surface topography along the length of fission yeast cells, including the presence of a septum scar from completion of a previous division with a cell pole undergoing new end polarized growth (Figure 1A, asterisk represents division septum scar). A height retrace image of the same cell (Figure 1B) provided cellular elevation data that was used to generate a 3-D reconstruction of the cell revealing that the peak height is 2.24 μm above the surface of the dish (Figure 1C). The height retrace image was also used for determining the optimal regions of the cell along the apex of cell curvature for taking force measurements (Figure 1B, red +) to determine the biophysical properties of the cell surface (Arfsten, Leupold, Bradtmoller, Kampen, & Kwade, 2010).

**Figure 1.**
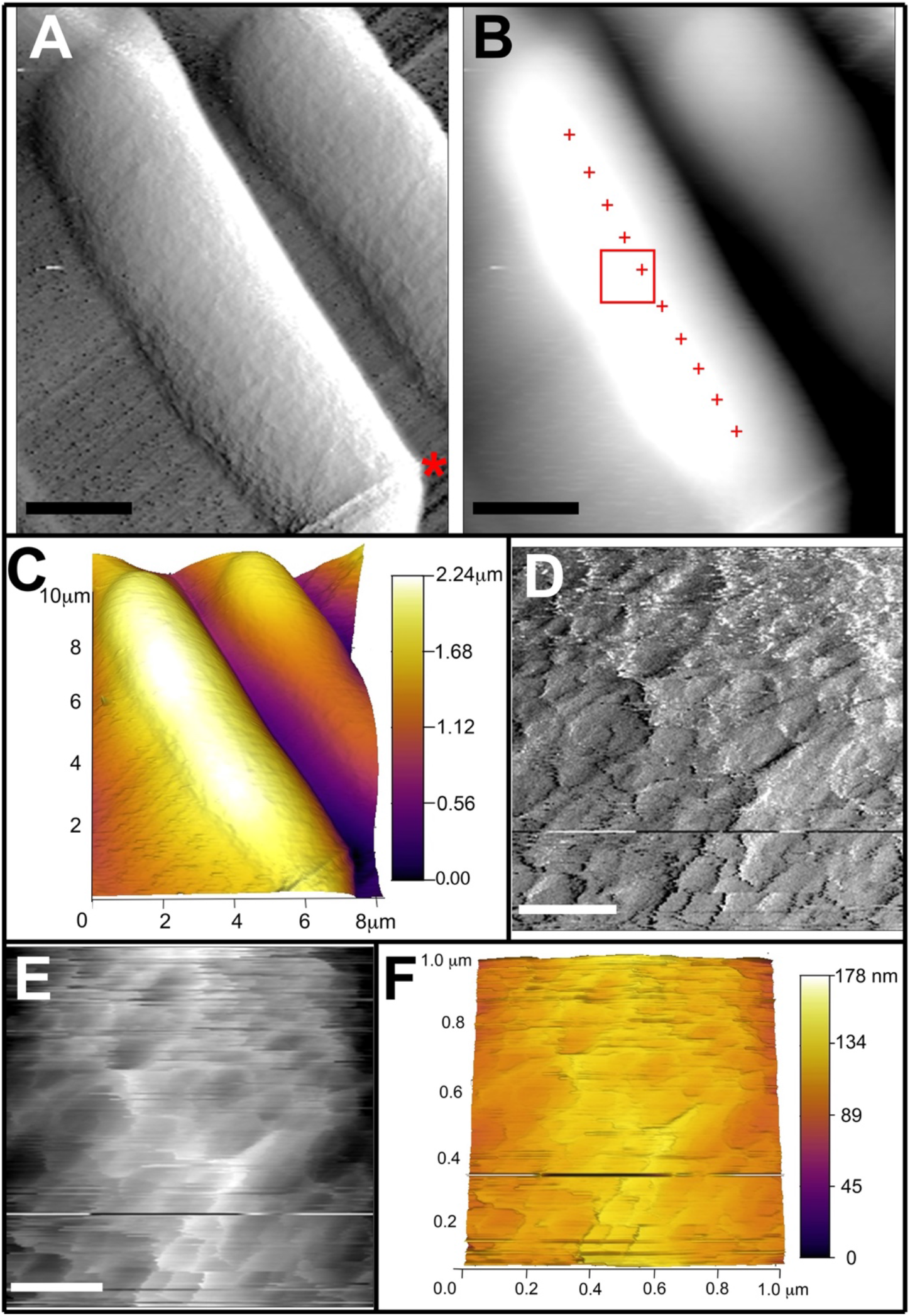
AFM topographical scanning of wildtype *S. pombe* cells. Atomic force microscopy (A) deflection retrace image and (B) height retrace image of wildtype cells acquired in contact scanning mode at 0.35 Hz at applied force of <1 nN. Scale bar for (A-B) is 2 μm. Asterisk in (A) represents division septum scar. Red boxed region in (B) indicates 1 μm × 1 μm area re-scanned for higher resolution shown in (D-F). Red ‘+’ indicate points sampled for force extension curves. (C) 3-D topography image of wildtype cell with heat map height scale ranging from 0 - 2.24 μm. (D) Deflection retrace image and (E) height retrace image of 1 μm × 1 μm region of interest acquired in contact scanning mode showing high resolution wildtype cell surface topography. Scale bar for (D-E) is 0.25 μm. (F) 3-D topography image of 1 μm × 1 μm region from wildtype cell with heat map height scale from 0 - 178 nm.

To gain further structural insight into the fission yeast cell wall topography at high-level resolution, a 1 μm × 1 μm region of interest was designated for additional contact mode scanning (Figure 1B, red box). The deflection retrace image (Figure 1D) of this area reveals non-uniform ridges in the cell wall that could represent unequal deposition or accumulation of cell wall components. This visualization of surface topography achieved by AFM scanning of live cells without fixation is comparable to other high-resolution fixed imaging approaches such as transmission and scanning electron microscopy (M. Osumi et al., 1998; Sipiczki, 2016). Furthermore, these images indicate that this cell surface topography is not an artifact of fixation or air-drying, but instead represents native structures on the fission yeast cell surface. The height retrace image (Figure 1E) and 3-D reconstruction of the cell surface (Figure 1F) reveal that these ridges extend up to 40 ± 11 nm from the average surface level of the cell with a calculated surface roughness (root mean square deviation) of 14 ± 3 nm (Table 1). These are the first reported roughness calculations for fission yeast cells maintained in an aqueous, non-stressed environment during analysis, and they are significantly lower than previous AFM cell surface roughness calculations for *S. pombe* cells following dehydration (69.9 ± 5.5 nm) or exposure to osmotic, thermal, ethanol, or oxidative stress (Adya et al., 2006; Canetta et al., 2006; Canetta et al., 2009). The fission yeast surface roughness is greater than AFM values reported for *S. cerevisiae* (~1-2nm) or *C. albicans* (8.2 ± 2nm), indicating that the *S. pombe* cell surface is less smooth, perhaps due to differences in cell wall composition (Ahimou, Touhami, & Dufrene, 2003; Alsteens et al., 2008; Dague et al., 2010b; Hasim et al., 2017).

**Table 1:** *S. pombe* cell surface roughness and biophysical properties.

| strain and sampling location | roughness: rms deviation (nm) | highest peak (nm) | cell wall elasticity: Δdistance (nm) | cell wall elasticity: Δforce (nN) | cellular spring constant (mN/m) | Young's modulus (MPa) |
| --- | --- | --- | --- | --- | --- | --- |
| wildtype: cell body | $14 \pm 3$ | $40 \pm 11$ | $65 \pm 24$ | $1.0 \pm 0.3$ | $44 \pm 10$ | $1.3 \pm 0.5$ |
| *cdc25-22*: cell body | $13 \pm 4$ | $37 \pm 11$ | $79 \pm 17$ | $1.1 \pm 0.3$ | $40 \pm 7$ | $1.4 \pm 0.4$ |
| *cdc25-22*: cell pole | $8 \pm 3$ | $16 \pm 2$ | $156 \pm 26$ | $1.4 \pm 0.3$ | $17 \pm 4$ | $0.5 \pm 0.1$ |

### Determining biophysical properties of fission yeast cells from force extension curves

After scanning the topography of the fission yeast cell wall, sites along the surface were designated for generating force curves by probing the cell with the AFM cantilever tip to determine biophysical properties of wildtype *S. pombe* cells. For each cell evaluated, a minimum of 100 force curves were generated at 10 different points (spaced ~0.5 μm apart) along the length of the cell (Figure 1B, red +). The extension force curves consist of the approach of the cantilever toward the surface (Figure 2A; horizontal linear region between −200nm to 0nm), the initial contact between the surface and the cantilever tip at low applied force (Figure 2A; non-linear region beginning at 0nm), and the indentation of the surface at higher applied force (Figure 2A; linear region of the curve). The non-linear region of the extension curve indicates the elasticity (Δforce and Δdistance) of the cell wall as it is compressed by the cantilever, while the linear region of the extension curve is used to determine the cellular spring constant and is related to the cellular turgor pressure providing greater resistive force to the cantilever (Arfsten et al., 2010; Arnoldi et al., 2000; Volle et al., 2008). As expected, the non-linear region of the extension curves of wildtype cells (Δforce: 1.0 ± 0.3 nN; Δdistance: 65 ± 24 nm) was significantly greater than the non-linear region of the dish surface (Δforce: 0.6 ± 0.1 nN; Δdistance: 23 ± 12 nm) indicating that the wildtype cell wall is more compressible and elastic than the solid dish surface coated with CellTak (Figure 2; Table 1). Notably, prior ultrastructural electron microscopy imaging revealed the thickness of the outermost galactomannan layer of the fission yeast cell wall was approximately 60-70 nm (Cortes et al., 2012; Masako Osumi, Konomi, Sugawara, Takagi, & Baba, 2006; M. Osumi et al., 1998), which suggests that the initial indentation observed in the non-linear region of these AFM force extension curves could correlate to compression of this outermost galactomannan layer prior to encountering increased resistance from the inner α- and β-glucan layer and cellular turgor pressure.

**Figure 2.**
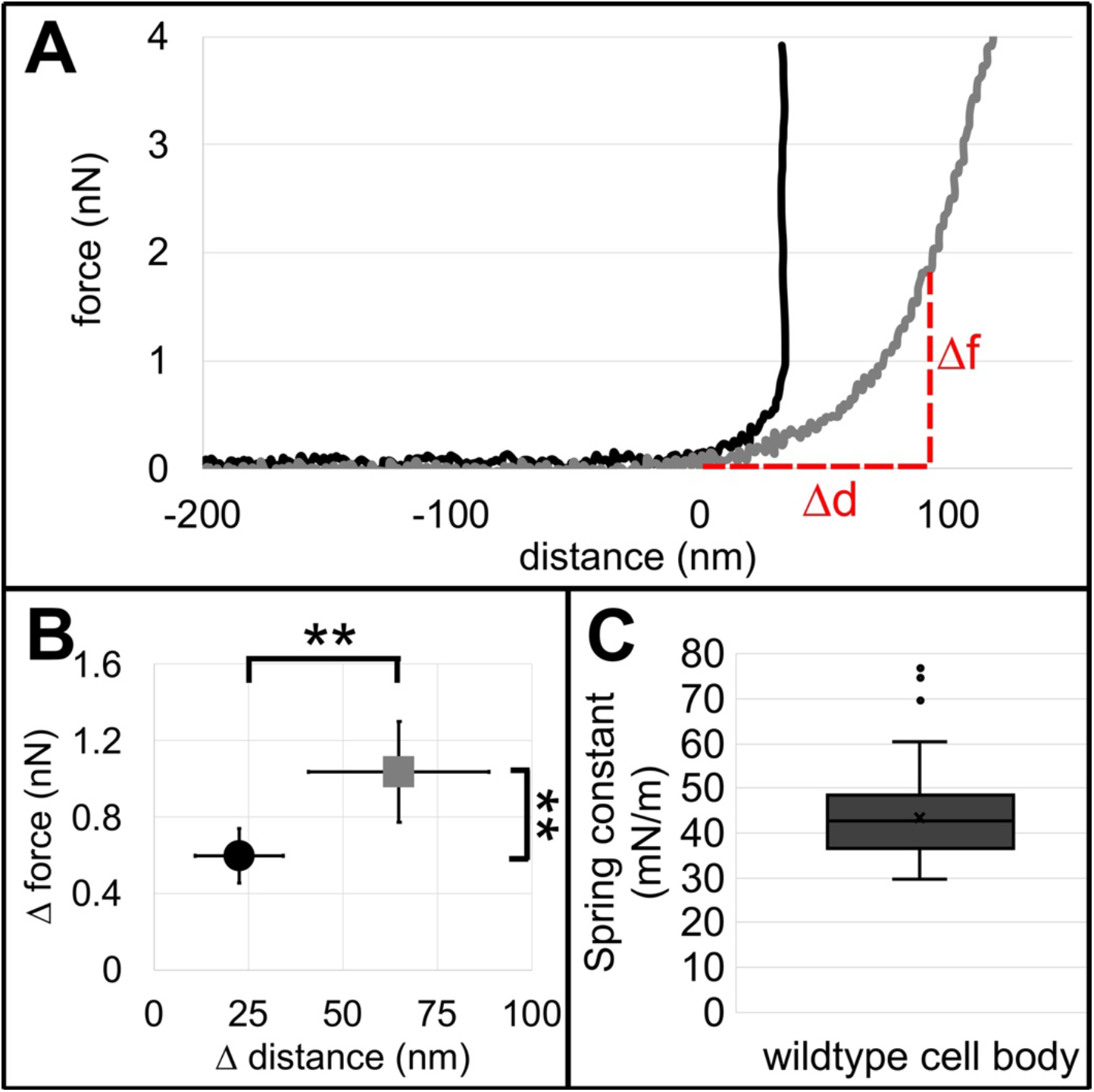
Characterization of wildtype *S. pombe* cell wall elasticity and cellular stiffness. (A) Representative force extension curves with 4 nN trigger point taken on the dish surface (black) or wildtype cells (gray) adhered to a plastic petri dish. The horizontal linear region of each curve extending from −200 nm to 0 nm reflects the cantilever approaching the surface, and the 0 nm point on the X-axis indicates the point of tip contact with the surface. The initial non-linear deflection of the force extension curve reflects the indentation of the cell wall upon initial contact, which is measured by Δf (change in force in nN) and Δd (change in distance in nm). The later linear deflection of the force extension curve indicates the cellular stiffness, which is related to the internal turgor pressure of the cell. (B) Plot of the elasticity (Δforce vs Δdistance) from the dish surface (black, circle) and wildtype *S. pombe* cell wall (gray, square) calculated from the nonlinear region of extension force curves. (error bars indicate s.d.; n > 100 force curves evaluated for each surface type on >10 different dishes or cells; two asterisks indicate p < 1e-10 determined by single factor ANOVA and paired student t-test). (C) Box-and-whisker plot of the wildtype cellular spring constant (*k*_cell_) calculated from the linear region of force extension curves acquired from points at the apex of cellular curvature along the length of the cell body (as determined by the AFM height retrace topography image). The boxed region indicates the upper and lower quartiles for each data set; the median is indicated by the horizontal line within the box; the mean is indicated by an ‘x’; whiskers extend to high and low data points; outliers are shown as individual data points. (n > 1000 force measurements from 10 wildtype cells).

The slope of the linear region of force extension curves were used to calculate spring constants for the dish and wildtype cell surface using Equation (1) and the experimentally determined spring constant of the cantilever. The wildtype cellular spring constant (*k*_cell_) of the *S. pombe* cell body was 44 ± 10 mN/m (Figure 2; Table 1). This *k*_cell_ is comparable to AFM measurements from other fungi including *Aspergillus nidulans* hyphae (*k*_cell_ of 29 ± 2 mN/m), and *Candida albicans* (*k*_cell_ of 51 ± 9 mN/m) (El-Kirat-Chatel et al., 2013; Zhao et al., 2005). AFM measurements of *Saccharomyces cerevisiae* spring constants vary widely (15 to 700 mN/m) depending on the region of the cell evaluated and experimental conditions utilized, but cells characterized using methodologies and conditions most similar to those in this study yielded a *k*_cell_ of 60 ± 25 mN/m (Arfsten et al., 2010; Pelling et al., 2004). Prior analyses of the *S. pombe* cell wall composition show significantly less glucosamine/chitin relative to *S. cerevisiae*, which could account for the lower *k*_cell_ observed in the fission yeast cell wall relative to budding yeast (Arellano, Cartagena-Lirola, Nasser Hajibagheri, Duran, & Henar Valdivieso, 2000; Dallies, Francois, & Paquet, 1998; Martin-Garcia, Duran, & Valdivieso, 2003; Sietsma & Wessels, 1990). Taken together, this evaluation of the fission yeast cell wall body by AFM reveals the previously uncharacterized biophysical properties of wildtype *S. pombe* cell wall elasticity and cellular stiffness.

### Biophysical characterization of the cell pole from force extension curves

Fission yeast cells utilize polarity signaling pathways, cytoskeletal alignment, and *de novo* glucan synthesis to promote polarized vegetative growth through extension at cell poles (Arellano et al., 2000; Chiou, Balasubramanian, & Lew, 2017; Martin et al., 2005; Mitchison & Nurse, 1985; Verde, Mata, & Nurse, 1995). The biophysical properties at these regions of active growth were evaluated to characterize differences in cell wall elasticity and/or cellular stiffness relative to the cell body where no extension or growth occurs. First, cells adhered to plastic dishes were imaged by AFM in contact scanning mode. Deflection and height retrace images were used to identify cells undergoing polarized tip growth through the characteristic presence of the residual septum from a prior division with a newly protruding pole (Figure 3A, asterisk). The area at the cell pole was then designated as the region of interest for force mapping (Figure 3B, red box) to determine whether there were sub-micrometer scale local differences in the nanomechanical cellular properties at the pole. Force mapping enabled the acquisition of 2,000 force extension curves within a 2.5 μm × 3.0 μm region (divided into a 40 × 50 grid). Spring constants (*k*_cell_) were calculated from each force extension curve and plotted within the 40 × 50 grid to determine variations in cellular stiffness at different regions within the cell pole (Figure 3D). *k*_cell_ values acquired from the area of the cell surface (as opposed to the dish) were plotted in a color-coded surface rendering for better visualization (Figure 3E), which revealed a gradient from low *k*_cell_ values (0-20 mN/m) at the apex of the cell pole to higher *k*_cell_ values (60-80 mN/m) at regions further from the tip and closer to the remnant of the septum.

**Figure 3.**
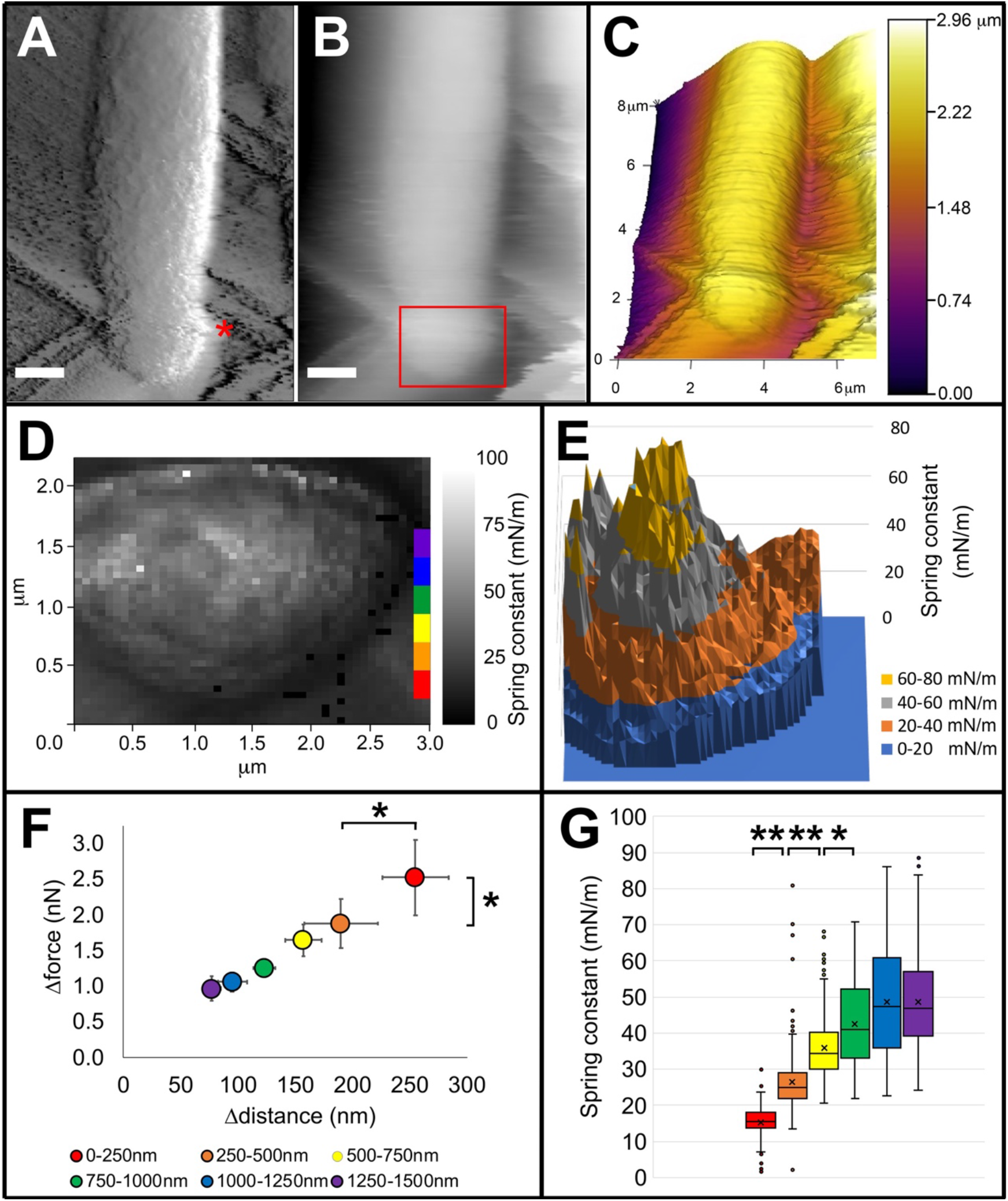
*S. pombe* cells adhered to dishes have decreased cellular stiffness at cell poles undergoing polarized growth. Atomic force microscopy (A) deflection retrace image and (B) height retrace image of wildtype cell acquired in contact scanning mode at 0.35 Hz at applied force of <1nN. Scale bar for (A-B) is 1 μm. Asterisk in (A) represents division septum scar. Red boxed region in (B) indicates 2.5 μm × 3 μm area used for generating force map shown in (D). (C) 3-D topography image of wildtype cell with heat map height scale ranging from 0 - 2.96 μm provided. (D) Force map of spring constant (*k*_cell_) values calculated from individual force extension curves acquired within the 2.5 μm × 3 μm region of interest (50 × 40 grid) at wildtype cell pole. Heat map of *k*_cell_ values ranging from 0-100 mN/nm provided. Red, orange, yellow, green, blue, and purple labels indicate locations of zones at cell pole evaluated in (F-G). (E) 3D color-coded surface rendering of *k*_cell_ values calculated from force extension curves on the wildtype cell pole force map shown in (D). (F-G) Measurements calculated from force extension curves were binned into 250nm zones (starting at the apex of the cell pole and moving progressively further away from the tip toward the septum scar) and averaged for each zone. (F) Plot of the surface elasticity (Δforce vs Δdistance) calculated from the non-linear portion of force extension curves from each zone of the cell pole. (error bars indicate s.d.; n > 20 force curves evaluated for each zone; one asterisk indicates p < 1e-5 determined by single factor ANOVA and paired student t-test). (G) *k*_cell_ calculated from force extension curves were binned and averaged for each zone. Box-and-whisker plots for *k*_cell_ from each zone. The boxed region indicates the upper and lower quartiles, the median is indicated by the horizontal line within the box, the mean is indicated by an ‘x’; whiskers extend to high and low data points; outliers are shown as individual data points. For (F-G) the most apical zone of the cell pole 0-250 nm (red); 250-500 nm (orange); 500-750 nm (yellow); 750 nm-1 μm (green); 1.0-1.25 μm (blue); 1.25-1.5 μm (purple). (n > 100 force curves evaluated for each zone; two asterisks indicate p < 1e-10, one asterisk indicates p < 1e-5 by single factor ANOVA and paired student t-test).

The cell pole was binned into 250 nm zones starting with the region closest to the pole apex and moving inward toward the septum remnant (Figure 3D, color bar inset), and the mean *k*_cell_ was calculated within each zone. The outermost 0-250 nm zone mean *k*_cell_ was significantly lower (15 ± 5 mN/m; Figure 3G, red, Table 2) than all other regions measured, indicating this region had decreased stiffness relative to the rest of the cell pole. The next outermost zone (250-500 nm; Figure 3G, orange, Table 2) *k*_cell_ increased to 26 ± 8 mN/m, which was significantly less than the stiffness of the 500-750 nm region *k*_cell_ (36 ± 9 mN/m; Figure 3G, yellow, Table 2). Mean *k*_cell_ values continued to increase in the 750 nm-1 μm zone (43 ± 12 mN/m; Figure 3G, green, Table 2) before leveling off with no significant difference between the 1-1.25 μm zone (49 ± 15 mN/m; Figure 3G, blue, Table 2), the 1.25-1.5 μm zone (49 ± 13 mN/m; Figure 3G, purple, Table 2). Similarly, elasticity in the outermost 0-250 nm region of the cell pole was greatest (Δforce: 2.5 ± 0.5nN; Δdistance: 255 ± 29 nm; Figure 3F, red, Table 2), and decreased in regions further from the cell tip apex (Figure 3F, Table 2). This suggests that regions of the cell pole where polarized growth occurs have the lowest cellular stiffness and greatest elasticity, and that stiffness increases further away from the apex of the cell pole. This is consistent with the need for greater flexibility and reduced rigidity at the pole as cellular extension occurs. These findings also reveal nanometer-scale differences within zones of the cell pole that could reflect progressive maturation of the cell wall as glucans are remodeled and cross-linked to provide increased stiffness along the cell shaft (de Medina-Redondo et al., 2010). Prior AFM characterization of *S. cerevisiae* lacking cell wall remodeling or cross-linking enzymes indicate decreased overall stiffness (Dague et al., 2010b). How homologous enzymes contribute to the biophysical properties of the fission yeast cell wall is an area of future investigation.

**Table 2:** *S. pombe* cell pole surface elasticity and stiffness.

| sampling location | cell wall elasticity: Δdistance (nm) | cell wall elasticity: Δforce (nN) | cellular spring constant (mN/m) |
| --- | --- | --- | --- |
| 0-250 nm | $255 \pm 29$ | $2.5 \pm 0.5$ | $15 \pm 5$ |
| 250-500 nm | $190 \pm 32$ | $1.9 \pm 0.3$ | $26 \pm 8$ |
| 500-750 nm | $157 \pm 16$ | $1.6 \pm 0.2$ | $36 \pm 9$ |
| 750nm -1μm | $123 \pm 9$ | $1.3 \pm 0.1$ | $43 \pm 12$ |
| 1-1.25μm | $95 \pm 13$ | $1.1 \pm 0.1$ | $49 \pm 15$ |
| 1.25-1.5μm | $77 \pm 8$ | $1.0 \pm 0.2$ | $49 \pm 13$ |

Notably, previous AFM studies of rod- and round-shaped microorganisms have shown variation of *k*_cell_ values dependent upon the distance at which measurements were taken relative to the apex of curvature of the cell, with a decreased *k*_cell_ at periphery zones, likely due to differences in the relative applied load acting on the cantilever tip (Arfsten et al., 2010; Gaboriaud, Parcha, Gee, Holden, & Strugnell, 2008). Therefore, we immobilized fission yeast cells vertically in 5 μm diameter well polycarbonate isopore membrane filters to enable force measurements at the cell pole apex of curvature. Wildtype *S. pombe* cells grow to an approximate length of 14 μm prior to dividing (Mitchison, 1957; Mitchison & Nurse, 1985; Moseley, Mayeux, Paoletti, & Nurse, 2009), which prevented AFM imaging of the cell pole due to the thickness of polycarbonate membrane filters (~22 μm). Therefore, we utilized *cdc25-22* cell cycle mutants that grow to approximately 21 μm prior to division due to delayed signaling for the onset of mitosis (Moseley et al., 2009; Nurse & Bissett, 1981; Thuriaux, Nurse, & Carter, 1978). This enabled the poles of these mutant cells to be imaged and probed while immobilized within the membrane filter wells (Figure 4A-B). Despite the genetic differences, the biophysical characteristics as measured by AFM for the cell body of *cdc25-22* cells were similar to the wildtype cell body with comparable force extension curves (Figure 5A; wildtype – gray, *cdc25-22* – red), Δforce (wildtype: 1.0 ± 0.3 nN; *cdc25-22*: 1.1 ± 0.3 nN), Δdistance (wildtype: 65 ± 24 nm, *cdc25-22*: 79 ± 17 nm) (Figure 5B), and spring constant (wildtype *k*_cell_: 44 ± 10 mN/m, *cdc25-22*: 40 ± 7 mN/m) (Figure 5C, Table 1).

**Figure 4.**
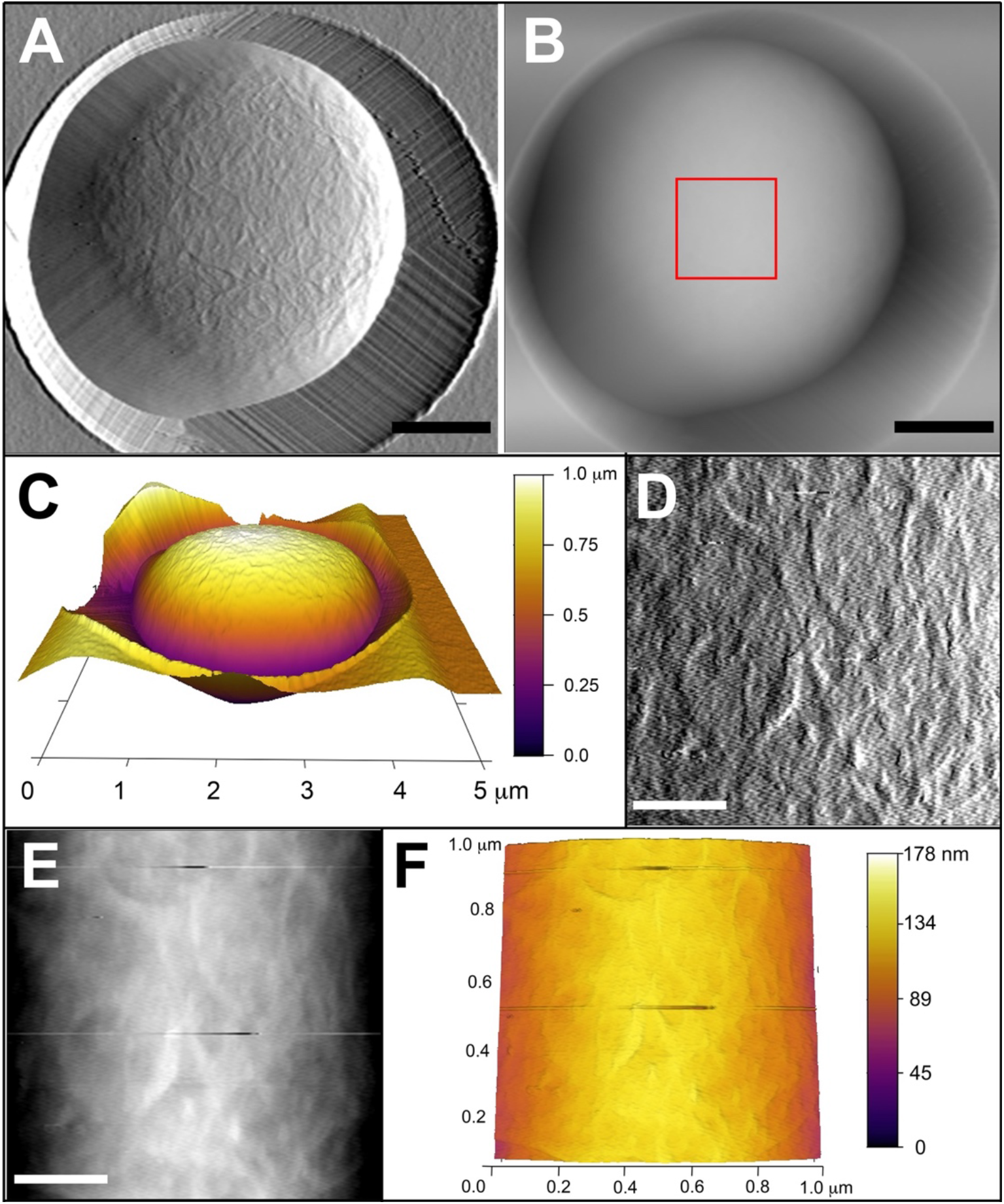
High resolution topographical scanning of *S. pombe* cell pole oriented vertically in a well. Atomic force microscopy (A) deflection retrace image and (B) height retrace image of *cdc25-22* cell immobilized within a 5 μm diameter pore in a polycarbonate membrane filter. Topographical scans acquired using contact scanning mode at 0.75 Hz and applied force of <1 nN. Scale bar for (A-B) is 1 μm. Red boxed region in (B) indicates 1 μm × 1 μm area re-scanned for higher resolution shown in (D-F). (C) 3-D topography image of *cdc25-22* cell within 5μm membrane pore with heat map height scale ranging from 0-1.0 μm. (D) Deflection retrace image and (E) height retrace image of 1 μm × 1 μm region acquired in contact scanning mode showing high resolution cell pole topography. Scale bar for (D-E) is 0.25 μm. (F) 3D topography image of 1 μm × 1 μm region from *cdc25-22* cell with heat map height scale from 0-178 nm.

**Figure 5.**
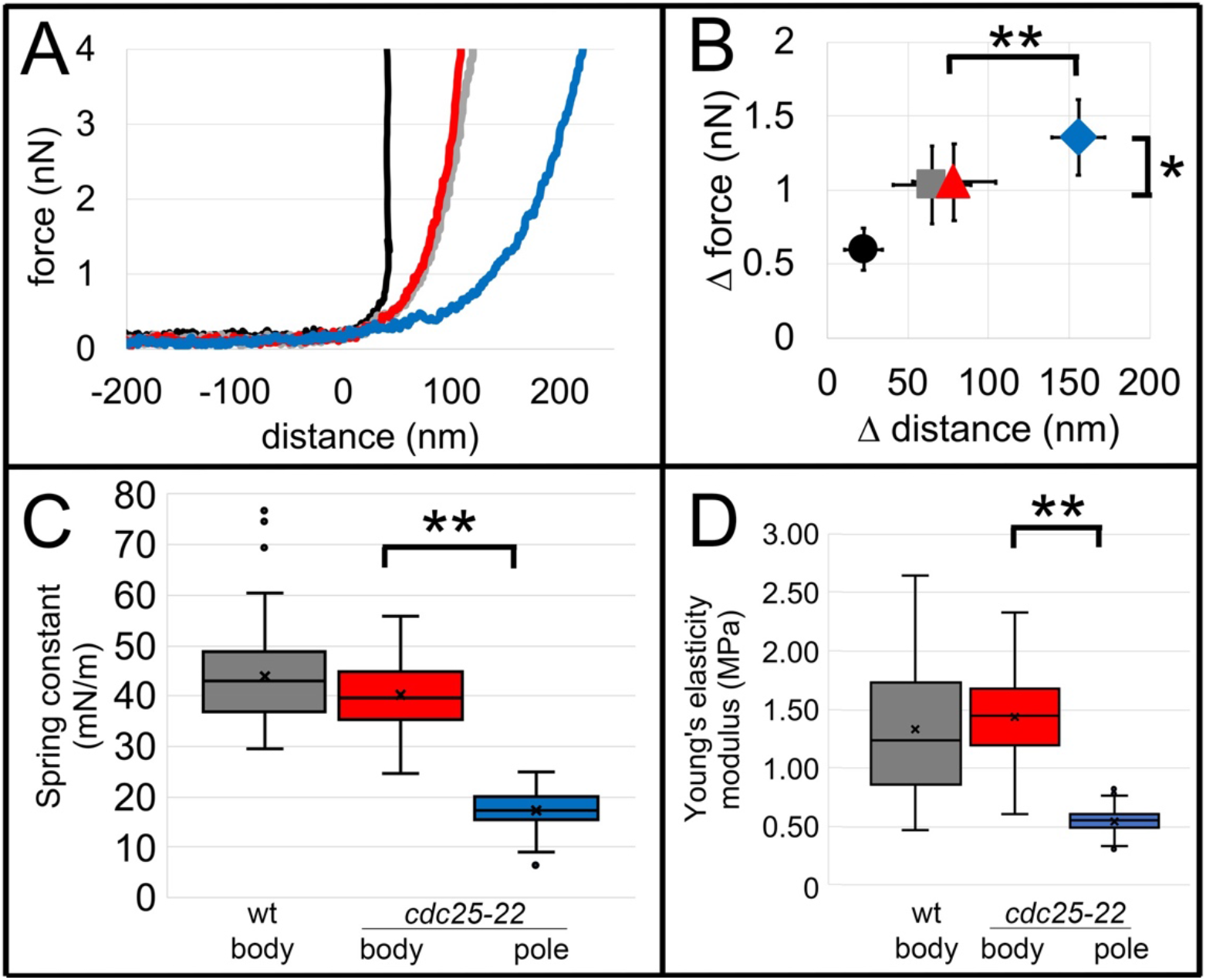
*S. pombe* cell poles have decreased cellular stiffness. (A) Representative force extension curves with 4 nN trigger point taken on the plastic dish surface (black), wildtype cell body (gray), *cdc25-22* cell body (red), or *cdc25-22* cell pole (blue). (B) Plot of the surface elasticity (Δforce vs Δdistance) from the plastic dish (black, circle), wildtype *S. pombe* cell body (gray, square), *cdc25-22* cell body (red, triangle), or *cdc25-22* cell pole (blue, diamond) calculated from the non-linear portion of force extension curves. (error bars indicate s.d.; n > 100 force curves evaluated for each surface type on >10 different dishes or cells per condition). (C) Box-and-whisker plot of *k*_cell_ calculated from the linear region of force extension curves from wildtype cell body (gray), *cdc25-22* cell body (red), or *cdc25-22* cell pole (blue). (n > 100 force measurements from 10 cells per condition). (D) Box-and-whisker plot of Young’s modulus of elasticity calculated by the Hertz model (using sample Poisson of 0.03) from cellular spring constants of wildtype cell body (gray), *cdc25-22* cell body (red), or *cdc25-22* cell pole (blue). For all box-and-whisker plots (C-D), the boxed region indicates the upper and lower quartiles for each data set; the median is indicated by the horizontal line within the box; the mean is indicated by an ‘x’; whiskers extend to high and low data points; outliers are shown as individual data points. (n > 100 force measurements from 10 cells per condition). For all graphs (B-D), two asterisks indicate p < 1e-10, one asterisk indicates p < 1e-5 determined by single factor ANOVA and student t-test.

Deflection retrace images from contact mode scanning of *cdc25-22* fission yeast cells immobilized within polycarbonate membrane filters provide cell pole surface topography (Figure 4A). The height retrace image of this same cell (Figure 4B) was used to generate a 3-D reconstruction of the cell pole inside the 5 μm well with the cell pole extending slightly above the polycarbonate membrane (Figure 4C). The height retrace was also utilized to designate a 1 μm × 1 μm region at the apex of the cell pole curvature for high-resolution scanning in contact mode (Figure 4B, red box). The deflection retrace image (Figure 4D) of this area has non-uniform cell wall ridges comparable in structure to those observed in the wildtype cell body (Figure 1D). However, the highest peak (16 ± 2 nm) and surface roughness calculated at the *cdc25-22* cell pole (8 ± 3 nm) was lower than either wildtype or *cdc25-22* cell body highest peak or roughness values (Table 1), indicating that the cell surface at the pole is flatter. This decreased elevation and roughness could be due to ongoing cell wall remodeling and restructuring at the pole during cellular extension and growth.

Force maps were generated from 900 force extension curves taken within 1 μm × 1 μm regions of interest (divided into a 30 × 30 grid) on the cell pole of immobilized *cdc25-22* cells in wells of polycarbonate membrane filters or the cell body of *cdc25-22* cells adhered to plastic dishes with CellTak adhesive. A representative force extension curve for the *cdc25-22* cell pole (Figure 5A) reveals a larger non-linear region with a significantly greater mean Δforce (1.4 ± 0.3 nN) and Δdistance (156 ± 26 nm) than the *cdc25-22* cell body (Δforce: 1.1 ± 0.3 nN; Δdistance: 79 ± 17 nm) or wildtype cell body (Δforce: 1.0 ± 0.3 nN; Δdistance: 65 ± 24 nm) (Figure 5B, Table 1). This suggests that the cell wall is more compressible and elastic at the pole than along the body where cellular growth and elongation are not occurring.

The slope of the linear region of force extension curves from *cdc25-22* cell poles was decreased relative to the wildtype or *cdc25-22* cell body (Figure 5A), and the *k*_cell_ of *cdc25-22* cell poles was significantly lower (17 ± 4 mN/m) than the *cdc25-22* cell body (40 ± 7 mN/m) or wildtype cell body (44 ± 10 mN/m) (Figure 5C). The *k*_cell_ for *cdc25-22* cell poles immobilized in wells of polycarbonate membrane filters was not significantly different from the spring constant observed in the outermost 250 nm bin of the force map acquired by probing wildtype cells adhered to a plastic dish with CellTak (Figure 3G, red). This suggests that the potential impact of differences in the relative applied load at the cell periphery during probing with the AFM cantilever was minimal in this study, and that wildtype cell poles exhibit a decrease in stiffness relative to the cell body.

Prior AFM studies in other microorganisms have applied Hertz modeling to calculate a Young’s modulus of cellular elasticity from local force extension curves and spring constants, though the applicability of this approach for microbial surfaces and how the resulting values can be compared to other measurements of elasticity moduli (such as those derived from whole cell compression experiments) is unclear and might require more complex finite element modeling (Arfsten et al., 2010; Gaboriaud & Dufrene, 2007; Mercade-Prieto, Thomas, & Zhang, 2013). However, to compare the biophysical measurements presented in this study with elasticity measurements from other fungi, we utilized the Hertz model (equation 2) to calculate Young’s moduli from the force extension curves of wildtype and *cdc25-22 S. pombe* cells. The wildtype cell body Young’s modulus was 1.3 ± 0.5 MPa, with the variation likely to due to local differences along the heterogenous surface of the cell (Figure 5D, Table 1). This value is consistent with estimates of the *S. pombe* internal turgor pressure that range from 1-1.5 MPa obtained by indirect methods of measurements or computational modeling (Abenza et al., 2015; Atilgan et al., 2015; Davi et al., 2018; Minc et al., 2009). Likewise, this value is similar to Young’s moduli (0.7–1.6 MPa) reported for *S. cerevisiae* cells when calculated by Hertz modeling of AFM force extension curves (Alsteens et al., 2008; Dague et al., 2010b; Pelling et al., 2004). The *cdc25-22* cell tip had a lower Young’s modulus of 0.5 ± 0.1 MPa, consistent with increased elasticity in this region (Figure 5D, Table 1). However, this does not reflect decreased turgor pressure at the cell pole, as the cell wall in this region is undergoing remodeling and maturation, which could contribute to the lower observed value. Further evaluation of fluctuations in the Young’s modulus of *S. pombe* cells in response to factors such as osmotic stress is necessary to provide additional insight into the utility of AFM force extension analysis as a tool for measuring the cellular turgor pressure in fission yeast.

This novel characterization of the relative biophysical properties of the *S. pombe* cell body and cell pole indicates that areas undergoing active cell growth and elongation have increased cell wall elasticity and decreased cellular stiffness. This study lays the groundwork for advancing high-resolution and quantitative analyses of the impact of various regulatory signaling pathways on cell wall elasticity during polarized cellular elongation and growth, characterizing components involved in polarized membrane trafficking during cell expansion, or evaluating the relative contributions of various cell wall synthases and cross-linking enzymes to establishing cell wall stiffness, all of which have historically been vital areas of research in the model organism *Schizosaccharomyces pombe*. Furthermore, this approach can be extended to studying polarized cell growth at the site of division during cytokinesis in fission yeast and determining how regulatory signaling pathways or cell wall synthase localization and activation contribute to new cell wall synthesis and ultimately septation and completion of division. Additionally, using AFM to calculate the cellular spring constant (*k*_cell_) and measure changes in cellular turgor pressure can be implemented in evaluating how fission yeast cells respond to environmental changes and provide new insights into the signaling pathways involved.

## ACKNOWLEDGEMENTS

The authors wish to thank Dr. Catherine Volle, Dr. Megan Nunez, Dr. Eric Darling, Dr. Qian Chen, and Dr. James Moseley for review and feedback on the manuscript. This work was supported by a National Science Foundation Major Research Instrumentation grant (number DBI1528288).

## GRAPHICAL ABSTRACT

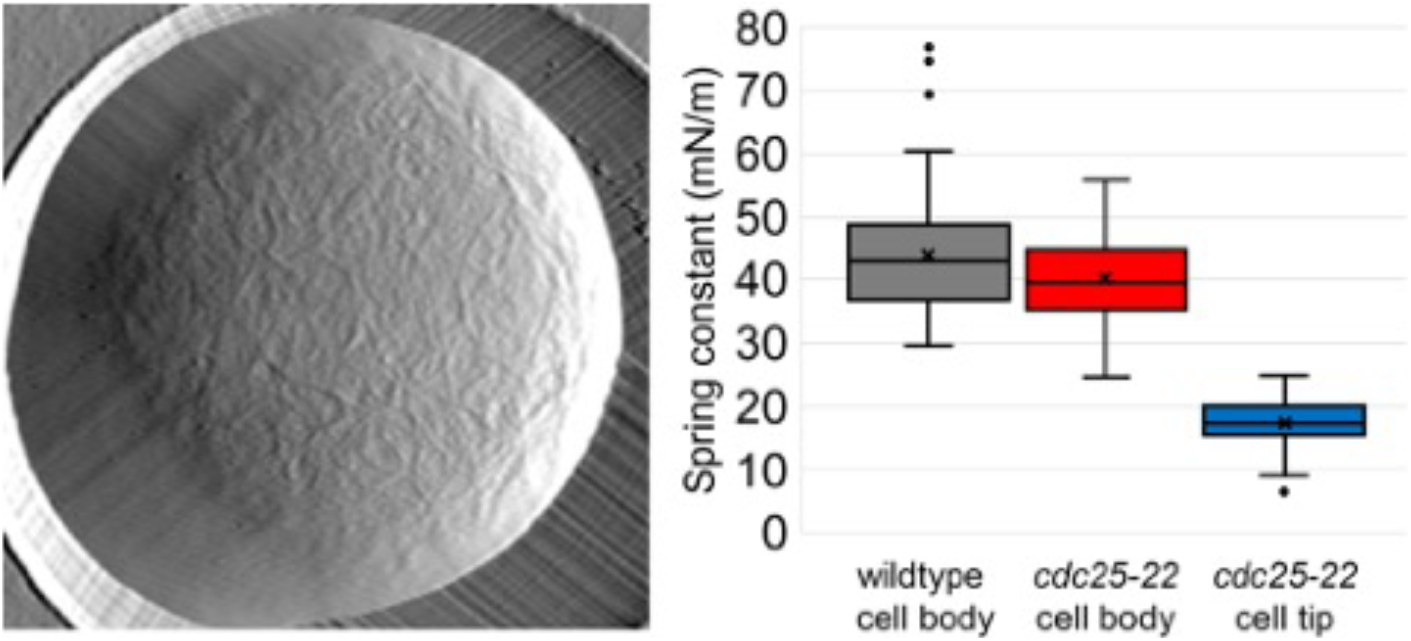

In this study, the authors utilize atomic force microscopy to evaluate the differential local surface topography and nanomechanical properties of fission yeast cells at regions of active growth relative to areas not undergoing expansion. They quantify decreased cellular stiffness at the poles, which could account for the plasticity necessary for polarized growth and expansion. These findings provide a foundation for future biophysical analyses of various cellular processes in fission yeast.

## SUPPORTING INFORMATION

**Supplemental Figure 1.**
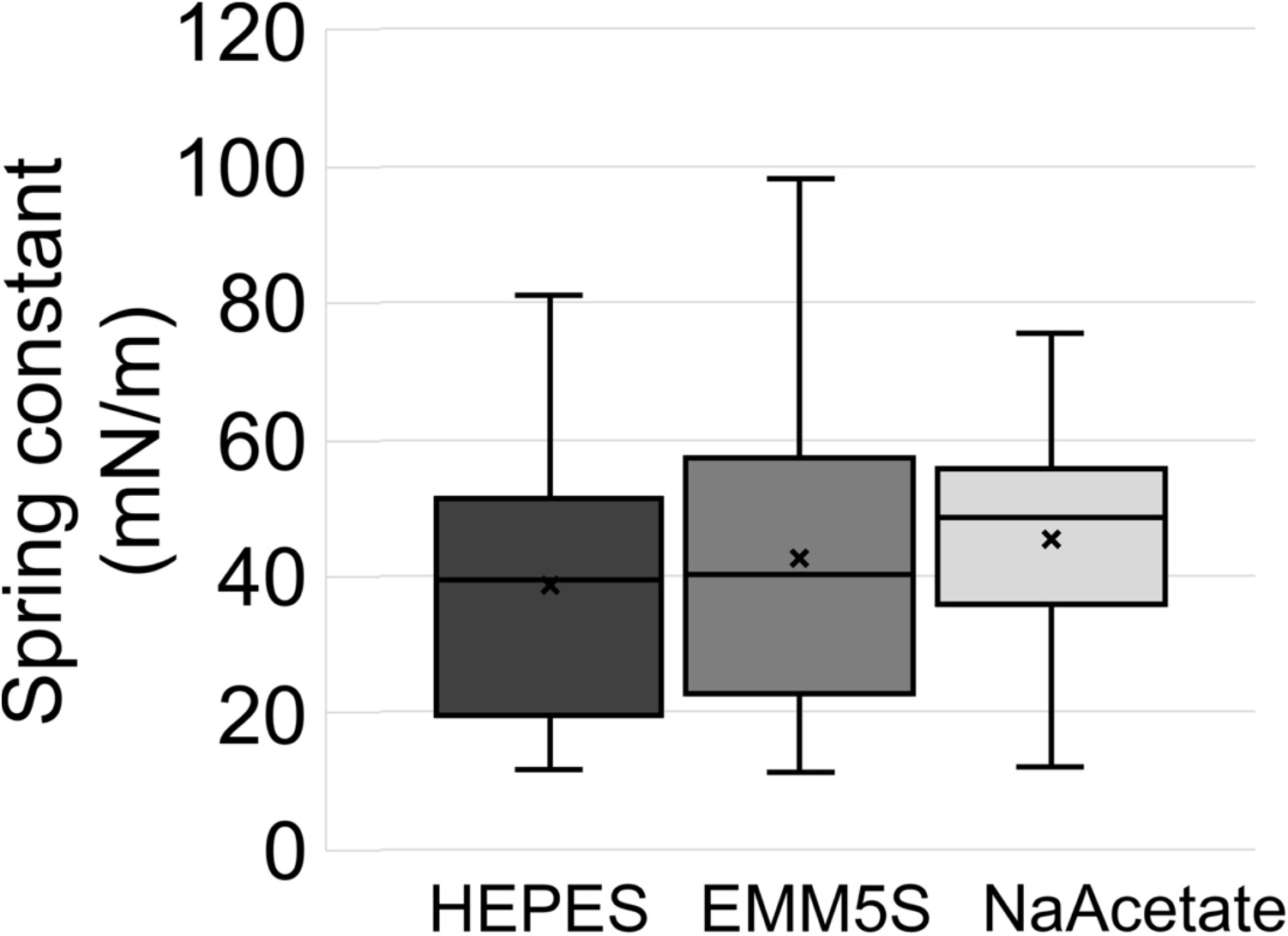
Comparison of *S. pombe* cellular stiffness in different imaging buffers. Wildtype *S. pombe* cells were incubated in 0.1 mM HEPES buffer, EMM5S fission yeast growth media, or 18 mM sodium acetate buffer during AFM imaging and probing. Box-and-whisker plots of the cellular spring constants were calculated from the linear region of force extension curves for each condition. The boxed region indicates the upper and lower quartiles for each data set; the median is indicated by the horizontal line within the box; the mean is indicated by an ‘x’; whiskers extend to high and low data points; outliers are shown as individual data points. (n > 500 force measurements from 5 cells under each condition).

**Supplemental Figure 2.**
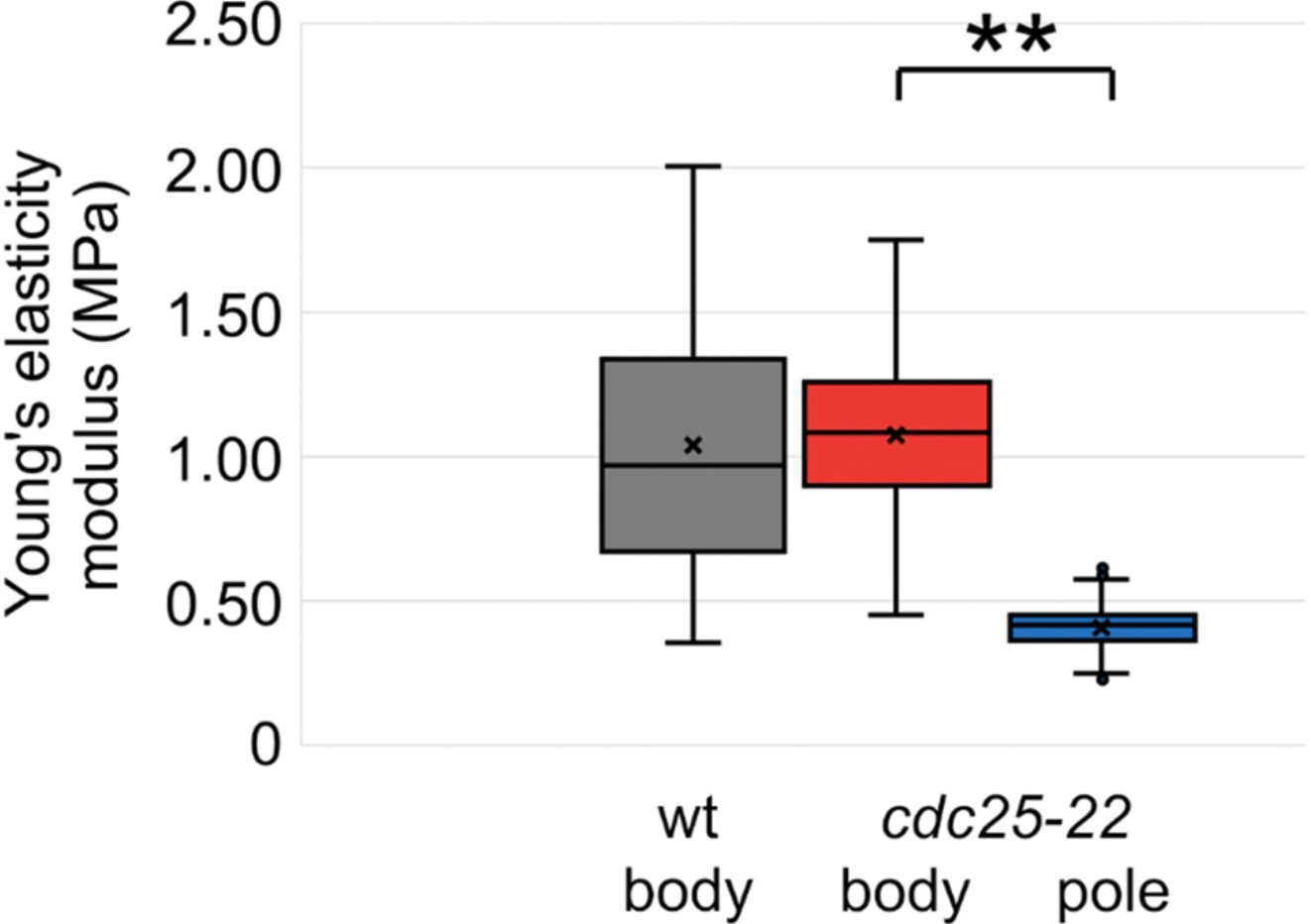
Calculation of Young’s modulus for *S. pombe* cells using Hertz modeling. Box-and-whisker plot of Young’s modulus of elasticity calculated by the Hertz model (using sample Poisson of 0.5) from cellular spring constants of wildtype cell body (gray), *cdc25-22* cell body (red), or *cdc25-22* cell pole (blue). The boxed region indicates the upper and lower quartiles for each data set; the median is indicated by the horizontal line within the box; the mean is indicated by an ‘x’; whiskers extend to high and low data points; outliers are shown as individual data points. (n > 100 force measurements from 10 cells per condition). Two asterisks indicate p < 1e-10, determined by single factor ANOVA and student t-test

**Supplemental Table 1:**
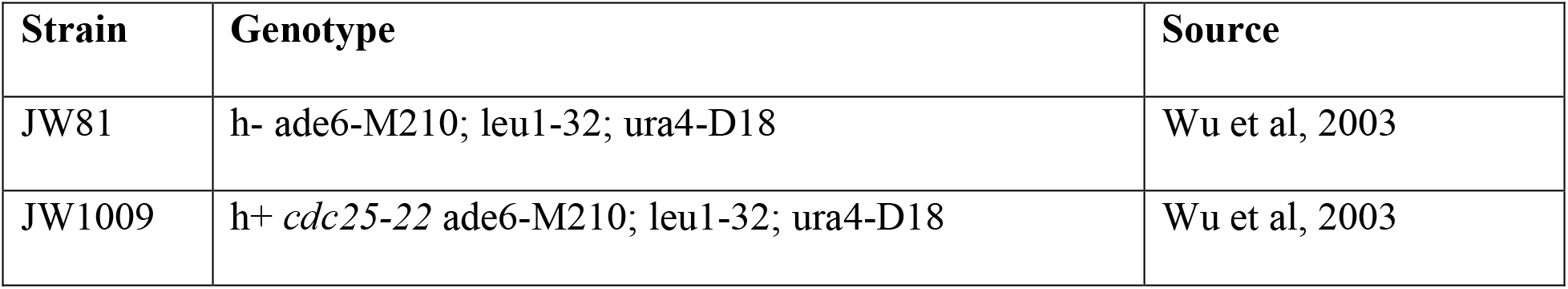
*S. pombe* strains used in this study.

